# Nitrogen availability and TOR signalling are important for preventing catastrophic mitosis in fission yeast

**DOI:** 10.1101/2023.12.25.573293

**Authors:** Viacheslav Zemlianski, Anna Marešová, Jarmila Princová, Roman Holič, Robert Häsler, Manuel José Ramos del Río, Laurane Lhoste, Maryia Zarechyntsava, Martin Převorovský

## Abstract

Mitosis is a critical stage in the cell cycle, controlled by a vast network of regulators responding to multiple internal and external factors. The fission yeast *Schizosaccharomyces pombe* may demonstrate catastrophic mitotic phenotypes due to mutations or drug treatments. One of the factors provoking catastrophic mitosis is a disturbed lipid metabolism, resulting from e.g. mutations in acetyl-CoA/biotin carboxylase (*cut6*), in fatty acid synthase (*fas2/lsd1*), or in the transcriptional regulator of lipid metabolism (*cbf11*) genes, as well as treatment with inhibitors of fatty acid synthesis. It was previously shown that mitotic fidelity in lipid metabolism mutants can be partially rescued by ammonium chloride. In this study we demonstrate that mitotic fidelity can be improved by multiple good nitrogen sources. Moreover, this rescue is not limited to lipid metabolism disturbances but also applies to a number of unrelated mitotic mutants. Interestingly, the rescue is not achieved by restoring the lipid metabolism state, but rather indirectly. We found that the TOR regulatory network plays a major role in mediating such rescue, highlighting a novel role for TOR in mitotic fidelity.

## INTRODUCTION

Mitosis is a critical stage in the cell cycle of any eukaryotic cell. Irregularities within this process may have far-reaching consequences, such as aneuploidy, mutations and/or cell death. These, in turn, can result in uncontrolled proliferation and tumour formation in metazoans [1], or cause decreased fitness in populations of unicellular organisms, such as yeasts [2]. Because of its significance, the entry into mitosis and progression through mitotic phases are tightly regulated by a network of redundant regulatory pathways, creating a fail-safe mechanism of control [3]. This network also integrates numerous internal and external factors that influence the decisions to proceed to the next phase. In yeasts, cell-cycle regulation is strongly affected by the availability of nutrients, such as carbon and nitrogen [4].

In contrast to the open mitosis of higher eukaryotes, many unicellular species undergo closed mitosis. The nuclear envelope (NE) remains intact during the whole process of a closed mitosis, even though substantial NE remodelling is required [5]. For example, in the fission yeast *Schizosaccharomyces pombe* the NE surface area needs to expand by 33% during mitosis to properly accommodate the elongating spindle and segregating chromosomes [6]. Notably, a number of abnormal mitotic phenotypes associated with closed mitosis have been described. These include: ‘cut’ - <u>c</u>ell <u>u</u>ntimely torn, a form of mitotic catastrophe where cytokinesis takes place before nuclear division has been properly resolved, and the mother nucleus gets transected by and trapped in the forming septum (Fig. 1 A) [7][8]; ‘lsd’ - large and <u>s</u>mall <u>d</u>aughters, a segregation error where the resulting daughter nuclei are of unequal sizes [9]; and nucleus displacement, which is similar to ‘cut’, but the forming septum misses the nucleus, resulting in one of the daughter cell being anucleate and the other one containing a diploid nucleus [10]. Typically, such events are lethal for at least one of the daughter cells. The ‘cut’ phenotype has been described for mutations in a range of mitosis-related genes, including the separase (*cut1*) and securin (*cut2*) [11], condensin (*cut3*) [12], anaphase-promoting complex (APC/C; *cut4*, *cut9*) [13], and others [14].

**Figure 1.**
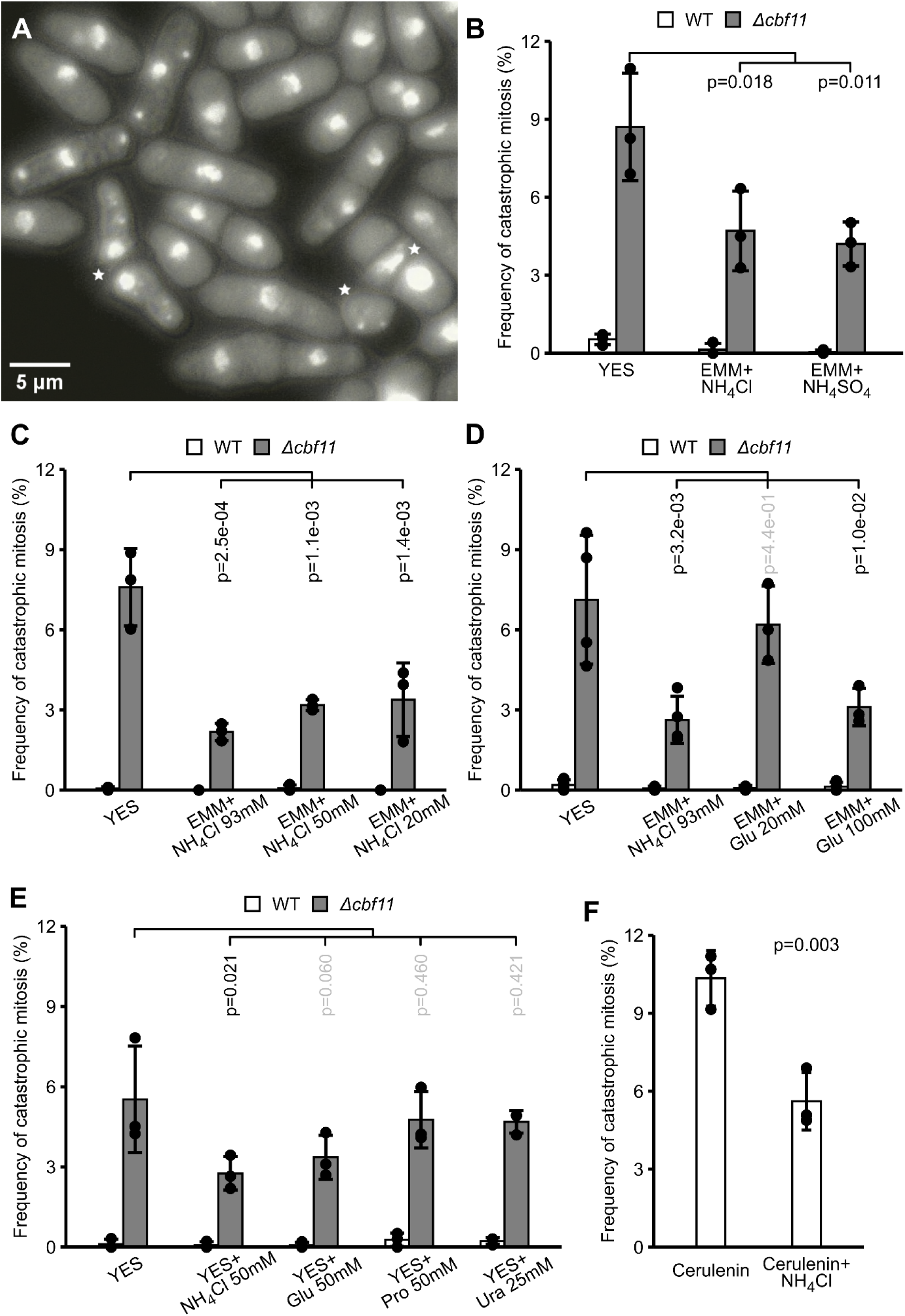
Addition of good nitrogen sources suppresses mitotic defects associated with perturbed lipid metabolism. Cells were grown to exponential phase in the indicated media, fixed, stained with DAPI and subjected to microscopy. **(A)** Examples of mitotic defects displayed by *Δcbf11* cells. Overlay of DAPI and DIC channels is shown; catastrophic mitoses are marked with asterisks. **(B, C)** Mitotic defects in *Δcbf11* cells are partially suppressed by ammonium supplementation at all concentrations tested. **(D, E)** Glutamate can also partially suppress mitotic defects in *Δcbf11* cells, but it is less potent than ammonium. Poor nitrogen sources (proline, uracil) do not suppress mitotic defects (uracil was used in 2x lower concentration because the molecule contains two atoms of nitrogen). **(F)** Ammonium can also partially suppress mitotic defects triggered by cerulenin treatment. Mean values ± SD from 3-4 independent experiments are shown in panels B-F. To determine statistical significance one-way Dunnet’s statistical test was applied for the multiple comparison with a control (B-E), one-way Student’s t-test was applied for the two-sample comparison (F).

Interestingly, perturbations of lipid metabolism can also cause mitotic catastrophe in *S. pombe* [15]. It was shown that mutations in acetyl-CoA/biotin carboxylase (*cut6*), fatty acid (FA) synthase (*fas2/lsd1*) [9], or the CSL transcriptional regulator of lipid metabolism (*cbf11*) genes [16], as well as treatment with FA synthesis inhibitors cerulenin [9] or Cutin-1 [17] lead to the ‘cut’ and/or ‘lsd’ phenotypes. It was assumed that decreased supply of precursors of membrane phospholipids (PL) leads to insufficient NE expansion during anaphase and mitotic failure [17]. However, we recently found additional factors contributing to decreased mitotic fidelity in cells with perturbed lipid metabolism, as *cbf11* and *cut6* mutants show altered cohesin dynamics and H3K9 modifications at the centromeric regions [18].

Nitrogen is an important macronutrient required for the synthesis of amino acids, nucleotides, and many other biomolecules, and the availability of nitrogen has a profound effect on the timing of entry into mitosis [19]. To coordinate growth and division with available resources the cell employs several nutrient-responsive regulatory pathways. In *S. pombe*, these include the protein kinase A (PKA), the stress-response MAP kinase Sty1, and, most prominently, the target of rapamycin (TOR) kinases [20]. Notably, similar mechanisms operate in mammalian cells as well [21]. There are two TOR complexes (TORCs) in *S. pombe*, each containing a different TOR kinase paralog. TORC1 (Tor2 kinase) is a major regulator of nitrogen metabolism and it stimulates growth and proliferation. On the other hand, TORC2 (Tor1 kinase) is involved in stress responses and maintenance of genome integrity. Importantly, there is crosstalk within the TOR network and the two TORCs operate in an antagonistic manner [22][23]. Moreover, apart from the quantity of available nitrogen, the exact chemical nature (quality) of a nitrogenous substance must be taken into account. Some, like ammonium and glutamate, are classified as ‘good’ sources that can be utilised easily and efficiently by the cells, while others, like proline, are ranked as ‘poor’ sources, in spite of them all containing equal amounts of nitrogen atoms per molecule [24][25].

Intriguingly, we previously demonstrated that the incidence of mitotic catastrophe in the *cbf11* and *cut6* lipid metabolism mutants is suppressed when cells are grown in the minimal defined EMM medium, which contains abundant ammonium chloride as the nitrogen source [16]. A similar rescue effect was achieved by growing mutant cells in the complex (and relatively nitrogen-poor) YES medium supplemented with ammonium chloride [26]. We now show that this rescue of mitotic fidelity is indeed nitrogen-dependent and is mediated by the TOR network. Unexpectedly, we found that nitrogen supplementation does not restore the expression of lipid-metabolism genes or lipid composition in *Δcbf11* cells, nor does it restore NE expansion during mitosis, indicating an indirect nature of the rescue. Moreover, we demonstrate that nitrogen supplementation can also rescue a range of other, non-lipid ‘cut’ mutants, suggesting a more general effect of nitrogen on mitotic fidelity. Our results highlight a previously unappreciated role of the TOR network in successful progression through mitosis.

## MATERIALS AND METHODS

### Strains, media and cultivation

Standard methods and media were used for the cultivation of *Schizosaccharomyces pombe* strains [27]. YES medium was prepared using Bacto Yeast Extract (BD Biosciences) and SP Supplements (Formedium). EMM medium was prepared using EMM Broth without Nitrogen (EMM-N; Formedium). Ammonium chloride (Sigma) or ammonium sulphate (Lachema/Chempol) were added to EMM-N to the final concentration of 93 mM, unless indicated otherwise. L-Glutamic acid monosodium hydrate, L-Proline or Uracil (Sigma) were added to the final concentrations from 20 to 100 mM, as required. *S. pombe* cell cultures were pre-grown for 8 hours at 32°C (25°C for temperature-sensitive strains) in 5 ml YES, then transferred to the medium with the appropriate supplements or stressors, diluted to OD_600_=0.005 (WT) or ∼0.03-0.05 (mutants) and incubated overnight to the early exponential phase (OD_600_=0.5) at 32°C or at appropriate semi-restrictive temperature. Cerulenin (Abcam) was added 2 hours before harvesting to the final concentration of 10 μg/ml. Rapamycin (Merck) was added at the time of starting the final culture to the final concentration of 0.3 μg/ml. Routine optical density (OD) measurements of liquid cell cultures were taken using the WPA CO 8000 Cell Density Meter (Biochrom). The strains used in the study are listed in Table S1. Deletion of *cbf11* was carried out using the pMP91 or pMP92 targeting plasmid, based on pCloneNAT1 or pCloneHYG1 respectively, as described [28] and confirmed by PCR. All other strains used in this study were constructed by standard genetic crosses. Plasmids and oligonucleotides used in this study are listed in Table S2 and Table S3 respectively.

### Microscopy

For nuclear staining, exponentially growing *S. pombe* cells were pelleted by centrifugation (1000 g, 3 min), fixed in 70% ethanol and stored at 4°C prior to imaging. Then cells were rehydrated in water and stained with 4′,6-diamidino-2-phenylindole (DAPI), 0.1 μg/ml final concentration. Samples were analysed using a Leica DM750 microscope with HC FL PLAN 100x/1.25 OIL objective. For each sample at least ten images (∼500-1500 cells total) were acquired. Frequency of catastrophic mitosis was counted manually using our standard scoring criteria [18].

For live-cell microscopy 1 μl of slightly resuspended cell pellet was applied on 2% agarose-YES media solidified in a 2 mm PDMS spacer [29], and covered with a coverslip. Slides were placed into an OKOlab environmental chamber set to 32°C. Time-lapses were acquired using Nikon Ti2 microscope with Plan Apo Lambda 60x Oil objective coupled with Hamamatsu ORCA-Flash4.0 camera in 16-bit, 2x2 binning mode. Snapshots were taken at 2 minute intervals. Images were acquired as Z-stacks, 5 to 9 slices with 0.3 μm step. Fluorophores were excited using CoolLED pE-4000 device. GFP fluorophore was filmed using 460 nm excitation at 21% light power, 100 (Cut11-GFP) or 50 ms (Hht2-GFP) exposure time and 510 nm emission filter. mCherry fluorophore was filmed using 580 nm excitation at 20% power, 200 nm exposure time and 590 nm emission filter.

Image analysis was performed using ImageJ software [30]. Z-stacks were processed as max-intensity projections. Time-lapses were corrected for photobleaching [31] and sample movement (“Linear Stack Alignment with SIFT”) [32].

### RNA-seq

Samples were prepared from 3 biological replicates. Cells were cultured to exponential phase, 10 mL were harvested (600 g, 2 min) and the cell pellet was flash-frozen with liquid nitrogen. Total RNA was isolated using hot acidic phenol method followed by phenol-chloroform extractions and precipitation [33]. Extracted RNA was treated with TURBO DNase (Thermo Fisher Scientific) and purified using RNeasy columns (Qiagen). RNA quality was assessed on Bioanalyzer 2100 (Agilent). Detailed sample preparation protocol is available at ArrayExpress database (see “Data availability”).

WT and *Δcbf11* samples were processed at the Institute of Clinical Molecular Biology, Christian-Albrechts-University Kiel, Germany. Libraries were prepared using the Illumina TruSeq stranded mRNA Library (poly-A), and sequenced in the pair-end mode on an Illumina NovaSeq 6000 instrument with the NovaSeq 6000 SP Reagent Kit with 100 cycles.

Samples of WT, *cbf11DBM, Δmga2, Δmga2Δcbf11, Pcut6MUT* and cerulenin treatment were processed at the Institute of Molecular Genetics, Czech Academy of Sciences, Prague, Czechia. The sequencing libraries were synthesised using KAPA mRNA HyperPrep Kit (Illumina platform) (Roche, KK8581) and analysed on an Illumina Nextseq 500 instrument using the NextSeq 500/550 High Output Kit v2.5 (75 Cycles) (Genetica, 20024906) with single-end, 75 bp, dual index 2x8 bp setup.

### RNA-seq data analysis

The reference *S. pombe* genome and annotation were downloaded from PomBase (2022-05-30; https://www.pombase.org) [34][35]. Read quality was checked using FastQC version 0.11.9 [36]. Adapters and low-quality sequences were removed using Trimmomatic 0.39 [37]. Clean reads were aligned to the *S. pombe* genome using HISAT2 2.2.1 [38] and SAMtools 1.13 [39]. Read coverage tracks were then computed and normalised to the respective mapped library sizes using deepTools 3.5.1 [40]. Mapped reads and coverage data were inspected visually in the IGV 2.9 browser [41]. Gene counts were generated using the GenomicAlignments package [42] in R/Bioconductor [43][44]. Datasets from different sequencing runs were normalised using RUVseq [45]. Differentially expressed genes (DEG) were detected using DESeq2 [46].

### Lipid analysis

Samples were prepared from 3 biological replicates. Cells were cultured to exponential phase in YES, 50 mL were harvested (1000 g, 3 min). Cell pellets were flash-frozen with liquid nitrogen and freeze-dried using HyperCOOL 3055 device (Gyrozen).

Total lipids were extracted using a method previously reported with minor modifications [47]. Briefly, dry cell weight (DCW) was determined gravimetrically prior to lipid extraction. Freeze-dried cells (approximately 10-15 mg) were mixed with 100 µL of ice-cold water followed by addition of 1 mL chloroform and methanol mixture (2:1, v/v). Cells were disrupted by FastPrep disintegrator (MP Biomedicals) with glass beads (diameter 0.4 mm, 3x40 s at the highest speed, with 5 min cooling on ice between cycles). Lipids were extracted by incubating in chloroform/methanol/water (1:2:0.8, v/v) and subsequently adjusting the mixture proportion to 2:2:1.8 (v/v) at room temperature. The organic phase containing the lipids was separated by centrifugation and dried under a stream of nitrogen. The resulting dry lipids were dissolved in 100 µL of chloroform and methanol mixture (2:1, v/v).

For fatty acid analysis total lipid extracts were transmethylated with 5% Na-OCH_3_ in methanol. Fatty acid methyl esters (FAME) were then extracted using *n*-hexane as described previously [48]. The analysis of FAME involved injecting 1 μL aliquots into a GC2010Plus gas chromatography (GC) apparatus (Shimadzu) equipped with a BPX70 capillary column (30 m × 0.25 mm × 0.25 µm, SGE Analytical Science) as described previously [47][49]. Identification of individual FAMEs was accomplished by comparing them with authentic standards of a C_4_−C_24_ FAME mixture (Supelco). The quantification of individual fatty acids was conducted using heptadecanoic acid methyl ester as an internal standard (Sigma Aldrich).

For the thin layer chromatography (TCL) an aliquot of lipid extract corresponding to 8 mg of DCW was applied to silica gel TLC plates (Merck) by a Linomat 5 semiautomatic sample applicator (Camag). Neutral lipids were separated by a two-step TLC solvent system using a method described previously (first step: petroleum ether/diethyl ether/acetic acid, 70:30:2; second step: petroleum ether and diethyl ether, 49:1) [50]. Individual lipid spots were visualised by charring the plates as previously reported [51]. Individual lipid spots were identified using lipid standards. Phospholipids were separated by the solvent system (chloroform/methanol/acetic acid/water, 75:45:3:1) as described previously [52].

## RESULTS

### The mitotic defects of *Δcbf11* cells can be rescued by a good nitrogen source

We have shown previously that mitotic defects in fission yeast lipid metabolism ‘cut’ mutants (Fig. 1 A) can be rescued by the addition of ammonium chloride to culture media [26]. As the first step toward understanding this phenomenon we compared the frequency of mitotic catastrophe in *Δcbf11* cells in the standard complex YES medium, and in the minimal defined EMM medium supplemented with ammonium chloride or ammonium sulphate. We found that the magnitude of the rescue effect was similar between the two ammonium salts (Fig. 1 B), indicating that the previously observed rescue had been caused by the ammonium, not the chloride.

Different nitrogen-containing compounds can be utilised by the cell with varying ease and efficiency (‘good’ vs ‘poor’ nitrogen sources) [24][25]. Therefore, we next tested several nitrogen-containing compounds for their ability to suppress mitotic defects in *Δcbf11* cells. We found that ammonium, a prototypical good nitrogen source, had the most profound effect, suppressing mitotic defects even at reduced concentrations (Fig. 1 C). Glutamate, another good nitrogen source, could also improve mitotic fidelity, albeit only at higher concentration (Fig. 1 D). On the other hand, the poor nitrogen sources proline and uracil did not suppress the mitotic defects of *Δcbf11* cells at the concentrations tested (Fig. 1 E). Thus, the magnitude of the rescue effect correlates with the preferability of a given compound as a nitrogen source to fission yeast cells [24][25].

Mitotic catastrophe can also be triggered by chemical inhibition of the fatty acid synthase (FAS). This is thought to be caused by insufficient supply of membrane building blocks for NE expansion during the anaphase of closed mitosis [6][17]. We found that the frequency of mitotic catastrophe in wild-type (WT) treated with the FAS inhibitor cerulenin [9] can also be partially rescued by ammonium supplementation (Fig. 1 F). This finding thus seems to be compatible with the hypothesis of membrane building block shortage being responsible for mitotic defects upon lipid metabolism disturbances [6][17].

### Ammonium does not suppress perturbations of lipid metabolism

To investigate whether ammonium supplementation suppresses mitotic defects via stimulating production of new membranes, we performed measurements of NE expansion during mitosis using live-cell microscopy. We used a fluorescently tagged nuclear pore protein, Cut11-GFP, as a NE marker [53]. When comparing *Δcbf11* and WT cells we found that the mutant cells have smaller NE cross-section, both at the start of mitosis and at the moment of maximum NE expansion. Importantly, we did not observe any differences between cultures grown with or without ammonium supplementation (Fig. 2). This indicates that the nitrogen-dependent rescue of mitotic fidelity is not achieved by enhancing the anaphase NE expansion. Indeed, we previously showed that ammonium supplementation did not correct the aberrant lipid droplet (LD) content of *Δcbf11* cells [26]. Also, we demonstrated that the defects in *Δcbf11* mitosis manifest already before anaphase [18].

**Figure 2.**
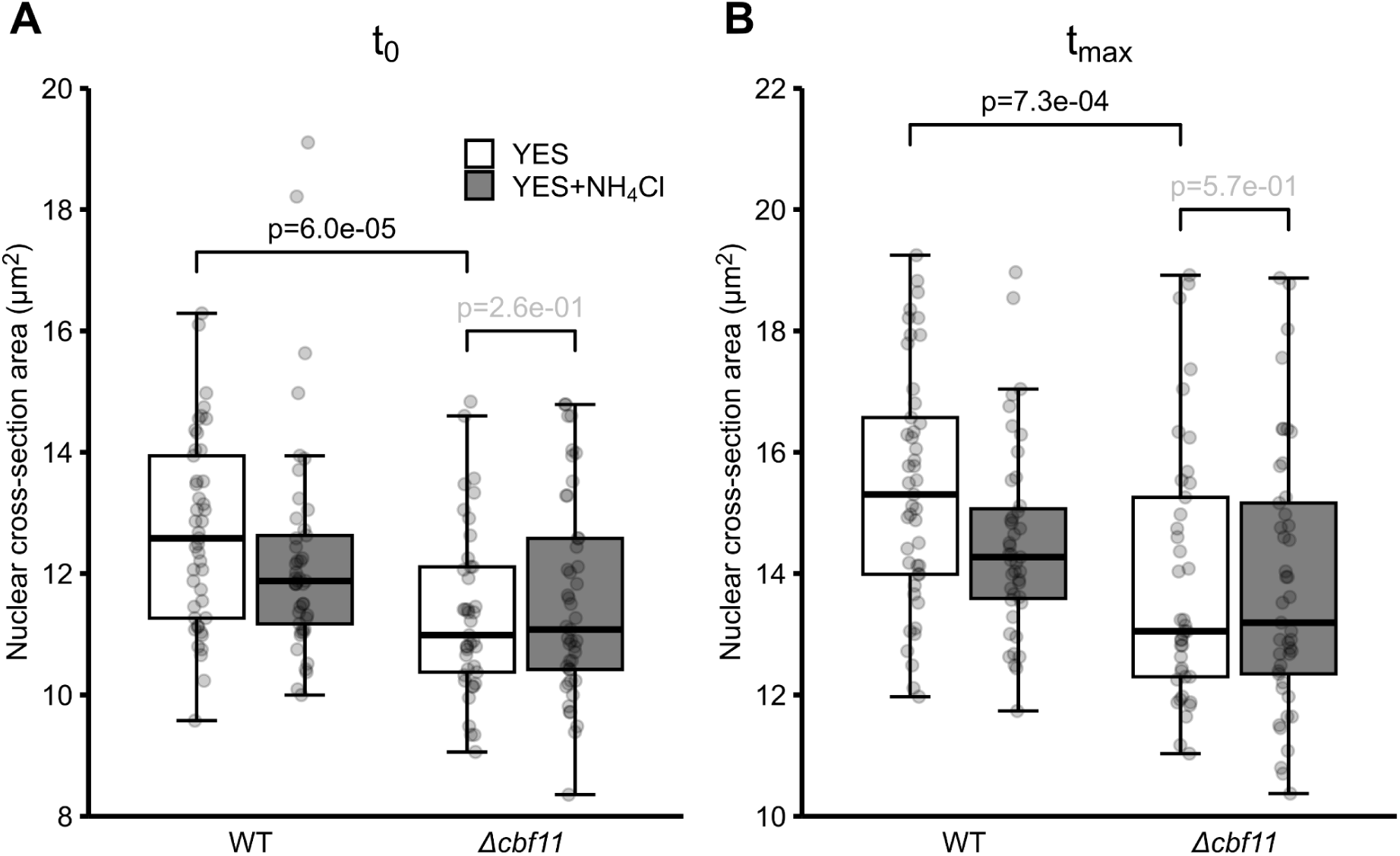
Small nuclear size in mitotic *Δcbf11* cells is not rescued by ammonium. WT and *Δcbf11* cells expressing the Cut11-GFP marker of NE were grown to exponential phase in the indicated media and subjected to time-lapse microscopy. The area of nuclear cross-section at the beginning **(A)** and end **(B)** of anaphase for 41-45 cells per condition is shown. Boxplots display medians (thick line), Q_1_ and Q_3_ quartiles (the box) and ± 1.5x interquartile ranges (whiskers). To determine statistical significance one-way Student’s t-test with Holm correction was applied.

To corroborate our new findings, we performed an RNA-seq analysis of a panel of lipid-metabolism mutants to assess the effect of nitrogen on the transcriptome. Our panel included the transcriptional regulators Cbf11 (*Δcbf11* and *cbf11DBM*, a mutant with abolished binding to DNA [54]), and Mga2 (*Δmga2*) [55], and the FA synthesis rate-limiting enzyme Cut6 (*Pcut6MUT* showing 50% reduction of *cut6* transcript levels [56]). As a control, we treated WT cells with the FA synthesis inhibitor cerulenin. We found that a number of lipid metabolism genes were downregulated in the *Δcbf11*, *Δmga2*, and *Δmga2 Δcbf11* double mutant, as well as in the *cbf11DBM* strain (Fig. 3). These genes include e.g. *cut6, fas1/2*, the acyl-coA desaturase *ole1*, and the long-chain-fatty-acid-CoA ligases *lcf1/2*. This is in agreement with the previously published studies of *Δcbf11* and *Δmga2* transcriptomes [16][55]. In contrast to the regulator mutants, targeted inhibition of FA synthesis (*Pcut6MUT*, cerulenin treatment) resulted in modestly increased expression of lipid metabolism genes, likely as a feedback reaction to FA shortage (Fig. 3). Importantly, lipid gene expression was not restored in regulator mutants grown in a medium supplemented with ammonium chloride (Fig. 3). This shows that the rescue of mitotic fidelity by a good nitrogen source is unlikely to be achieved by boosting the expression of lipid metabolism genes.

**Figure 3.**
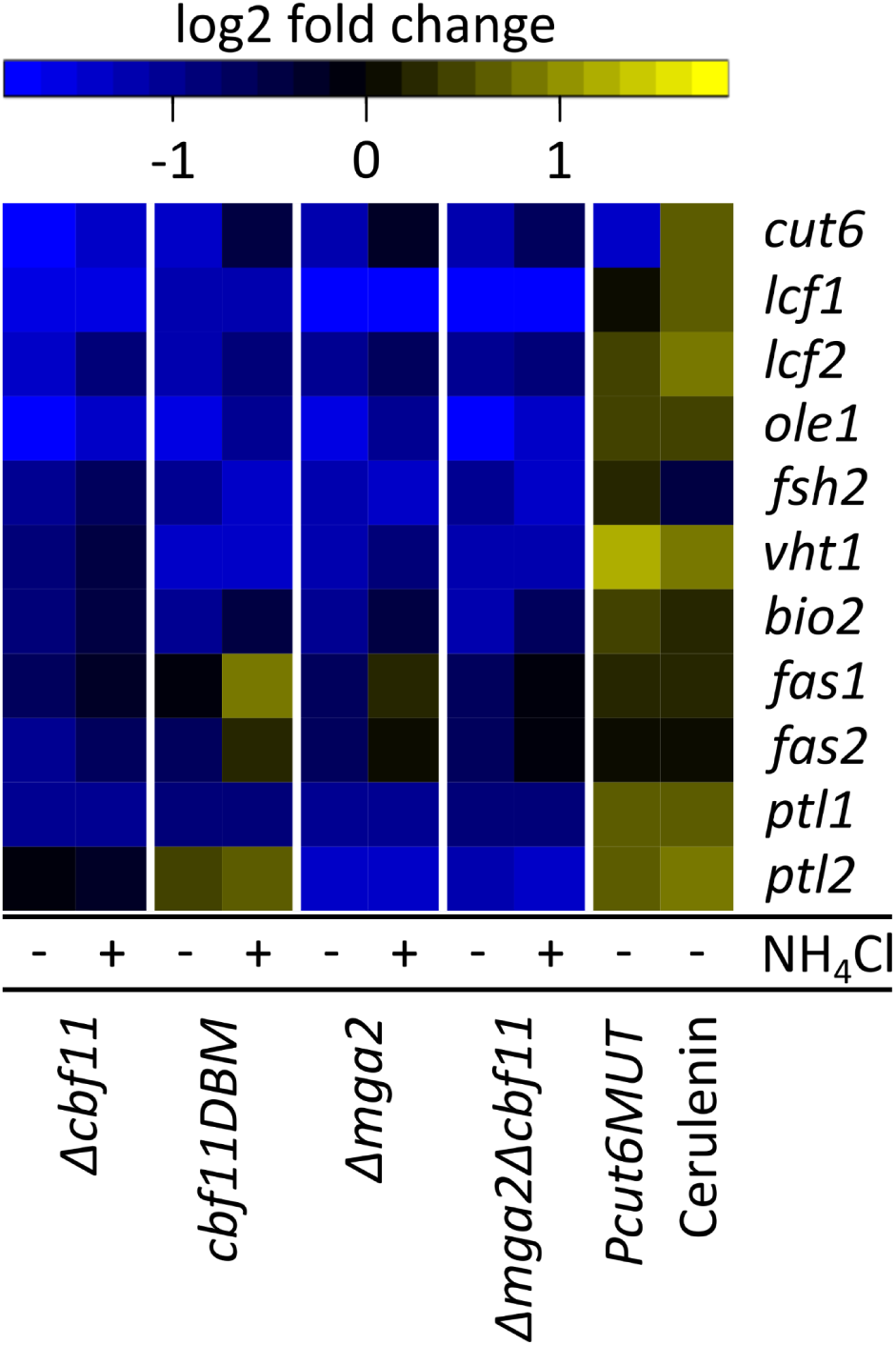
Decreased expression of lipid metabolism genes in *cbf11* and *mga2* mutants is not rescued by ammonium. RNA-seq results are shown as log2 of expression fold change values normalised to WT. Gene expression in the *Pcut6MUT* mutant and in WT treated with cerulenin are shown for comparison.

To further test for any potential effects of a good nitrogen source on the lipid metabolism in mitotic mutants, we analysed the total lipid content of *Δcbf11* and WT cells using thin layer chromatography (TLC) and FA composition by gas chromatography. The results showed that the total amount of FA is decreased in *Δcbf11* cells (Fig. 4 A), which correlates well with our previous data on decreased number of storage LDs in this strain [16]. Also, total FA saturation level is higher in *Δcbf11* cells compared to WT (Fig. 4 B), which may be caused by the decreased expression of the *ole1* desaturase gene (Fig. 3). This tendency towards higher FA saturation is also visible at the level of individual FA species (Fig. 4 C; compare C18:1 with C18:0 and C16:0). Finally, the TLC analysis revealed that the content of squalene and sterol esters is markedly increased in *Δcbf11* cells (Fig. 4 D, Fig. S1), suggesting failed coordination between the sterol and the triglyceride lipid metabolism pathways [57]. Notably, none of the detected changes in the *Δcbf11* FA and lipid composition were reversed by ammonium supplementation (Fig. 4, Fig. S1, Fig. S2). Taken together, our results strongly suggest that a good nitrogen source does not rectify the disturbed lipid metabolism in *Δcbf11* cells, and therefore it likely brings about the mitotic rescue in a different, indirect way.

**Figure 4.**
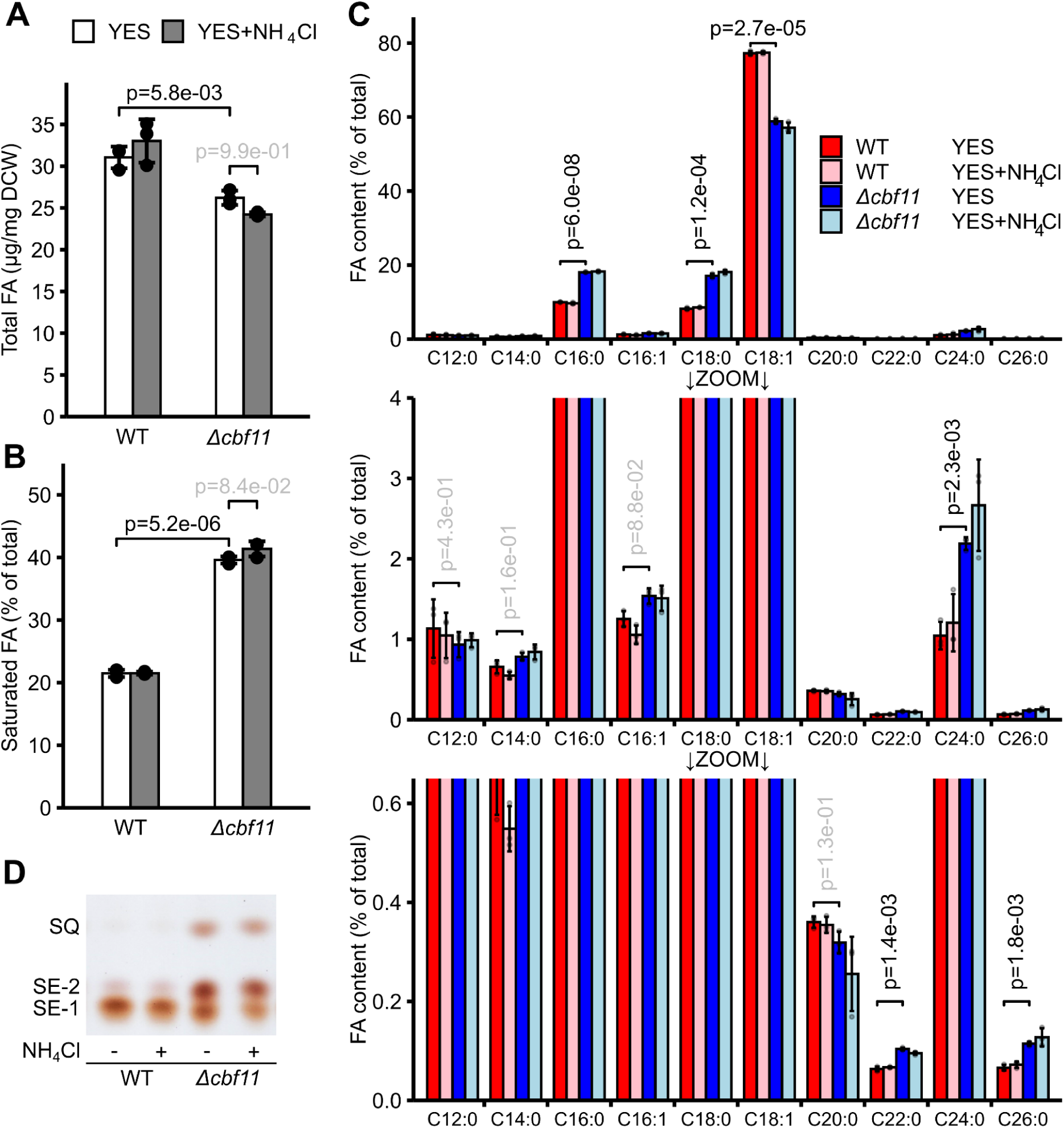
Lack of *cbf11* leads to pronounced changes in FA and lipid composition which are not affected by ammonium supplementation. WT and *Δcbf11* cells were grown to exponential phase in the indicated media, and their lipid composition was analysed. (A) *Δcbf11* cells have lower FA content per unit of dry cell weight compared to WT. **(B)** Degree of FA saturation is higher in *Δcbf11* cells. **(C)** Abundance of selected FA species in total lipid samples. Two different y-axis scales are shown to better visualise the full range of FA abundances. Mean values ± SD from 3 independent experiments are shown in panels A-C. To determine statistical significance one-way (A) or two-way (B, C) Student’s t-test with Holm correction was applied. **(D)** Thin layer chromatography (TLC) analysis of neutral lipids. Only a section of the TLC plate is shown (see Fig. S1 for the full TLC plate). SQ - squalene; SE-1, SE-2 - sterylester species.

### Ammonium has a general positive effect on mitotic fidelity

We previously showed that the duration of mitotic phases in *Δcbf11* cells is typically longer and more variable compared to WT cells. The delays first manifest well before the anaphase and they accumulate during the whole mitotic duration [18]. We now investigated whether mitotic timing is affected by a good nitrogen source. We performed live-cell microscopy of strains with fluorescently tagged histone H3 (Hht2-GFP) and alpha-tubulin (mCherry-Atb2) to visualise the chromatin and mitotic spindle, respectively (Fig. 5 A). We found that ammonium reduced the overall length of mitosis (Fig. 5 B), and this effect manifested both during the prophase and metaphase (Fig. 5 C), as well as anaphase stages (Fig. 5 D) in *Δcbf11* cells, bringing them close to WT values. Also, mitotic timing became more uniform among individual *Δcbf11* cells (Fig. 5). Note that telophase duration was not included in the overall mitosis length due to technical limitations of our experimental setup.

**Figure 5.**
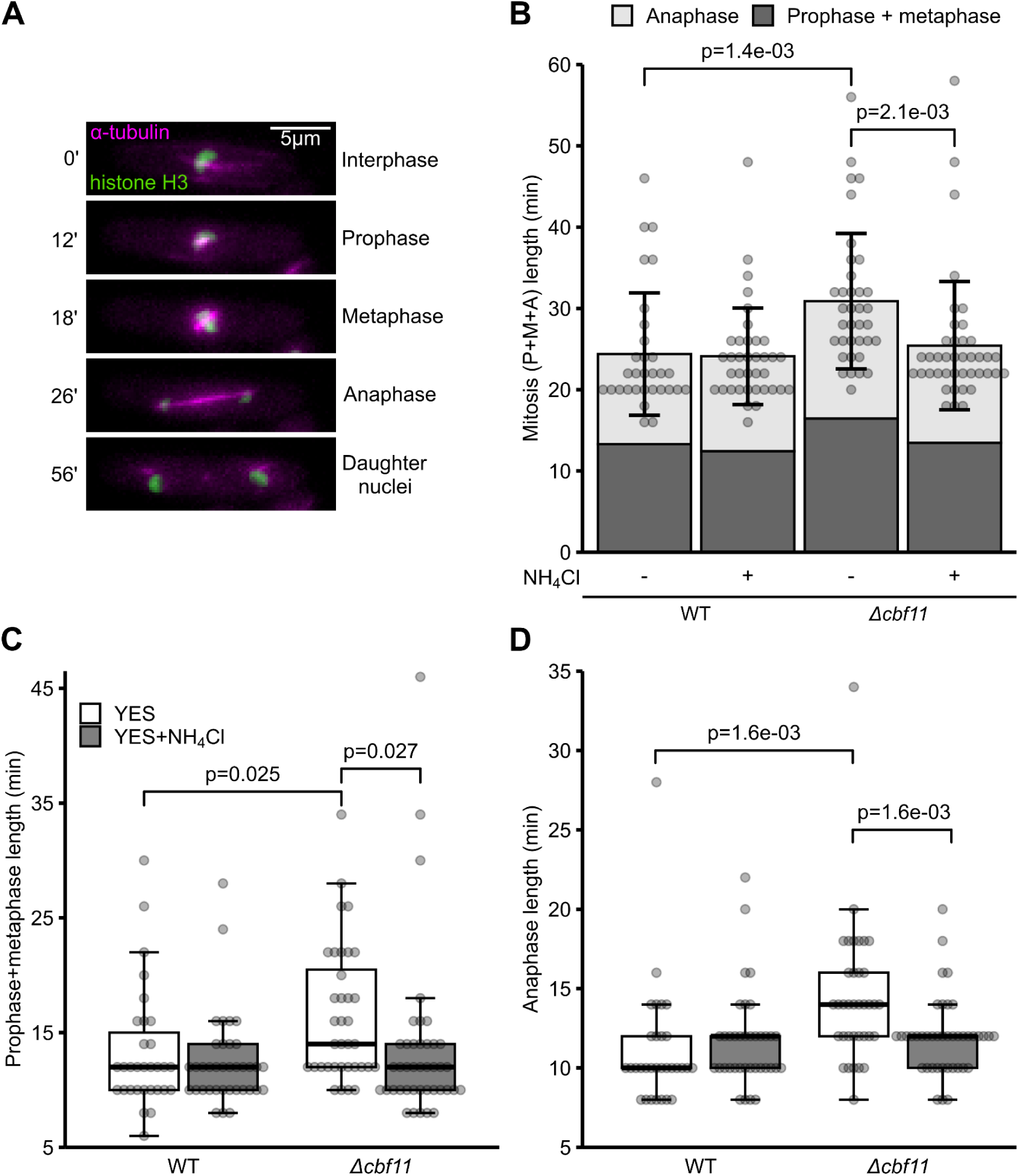
Ammonium rescues prolonged mitotic duration in *Δcbf11* cells. Examples of mitotic phases as they are observed under the microscope. Green - chromatin (Hht2-GFP), magenta - microtubules (mCherry-Atb2) **(A)**. The duration of the whole mitosis (prophase, metaphase and anaphase combined) **(B)**, as well as prophase+metaphase **(C)** or anaphase **(D)** separately are all prolonged in cells lacking *cbf11*. This aberrant timing does not occur in the presence of ammonium. Data for 31-41 cells per condition are shown. Barplot (B) displays mean values ± SD together with individual data points. Boxplots (C, D) display medians (thick line), Q_1_ and Q_3_ quartiles (the box) and ± 1.5x interquartile ranges (whiskers), together with individual data points. To determine statistical significance one-way Student’s t-test with Holm correction was applied. Only cells which successfully completed mitosis were analysed.

Our results so far indicated that nitrogen may have a more general effect on mitotic fidelity. To test this hypothesis, we determined the effect of ammonium on a diverse panel of mutants, none of them being related to lipid metabolism, which are prone to develop the ‘cut’ phenotype with varying severity. Remarkably, ammonium supplementation significantly decreased the frequency of catastrophic mitotic events in many, but not all, of those mutants (Fig. 6 A-C). The group of rescued ‘cut’ mutants consisted of condensin (*cut3*), cohesin (*psm3*), and the SMC5/6 complex (*smc6*, *nse3*), a nuclear proteasome tethering factor (*cut8*) and a nuclear import factor (*cut15*). On the other hand, the mutants without significant rescue were separase (*cut1*) and securin (*cut2*), and APC/C subunits (*cut4*, *cut9*). It should be noted though that the magnitude of the rescue varies between the ammonium-responsive mutants, and we suggest possible explanations for this behaviour in the Discussion. Taken together, the ammonium-mediated rescue of mitotic defects is not limited to cells with perturbed lipid metabolism, and seems to operate early in the cell cycle, prior to anaphase.

**Figure 6.**
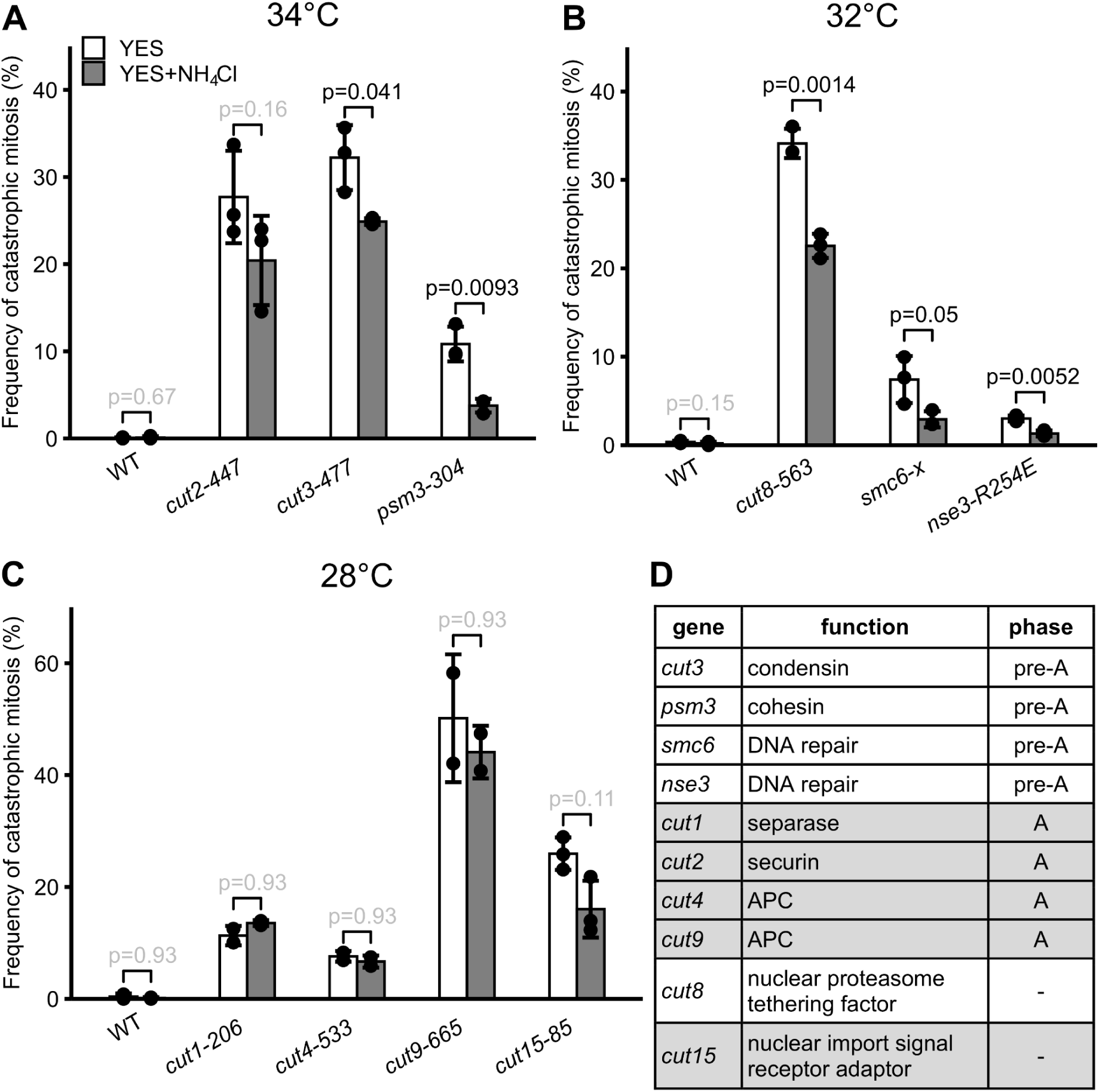
Ammonium supplementation rescues a range of cut mutants. **(A-C)** Cells were grown to exponential phase in the indicated media, fixed, stained with DAPI and subjected to microscopy. Cultivation temperatures were chosen to provide semi-restrictive conditions for temperature-sensitive strains. Mean values ± SD from 2-3 independent experiments are shown. To determine statistical significance one-way Student’s t-test with Holm correction was applied. **(D)** Mutants responsive to ammonium supplementation tend to be associated with pre-anaphase (“pre-A”) rather than anaphase (“A”). Grey background - no rescue.

### The nitrogen-dependent rescue of mitotic fidelity is mediated by TOR

The evolutionarily conserved TOR signalling network is a major hub for controlling nitrogen sensing and utilisation. Therefore, we tested whether and how TOR was involved in the nitrogen-dependent improvement of mitotic fidelity. First, we suppressed the activity of TOR by treating cells with the TOR inhibitor rapamycin [58]. Notably, we found that rapamycin rescued mitotic fidelity in *Δcbf11* cells to an extent similar to that of ammonium. Additionally, a combined treatment showed that the effects of rapamycin and ammonium were non-additive (Fig. 7 A). These results strongly indicate that both chemicals affect the same target pathway or process.

**Figure 7.**
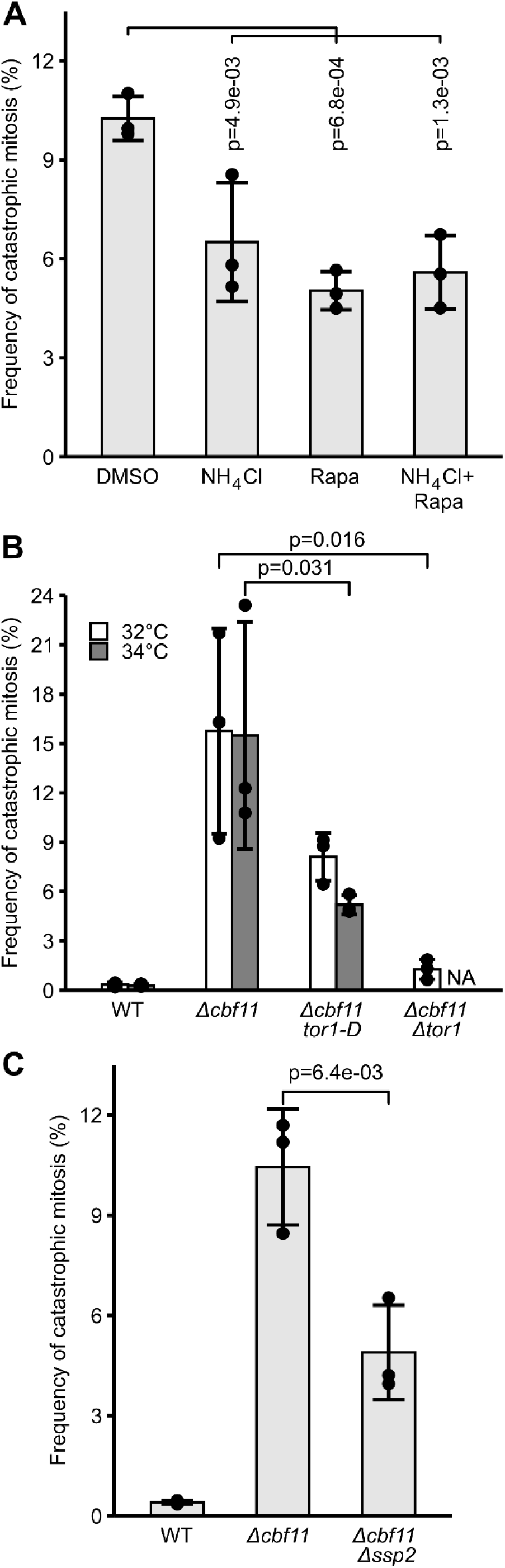
The TOR network is critical for the ammonium-mediated rescue of *Δcbf11* mitotic defects. **(A)** Rapamycin treatment partially suppresses the mitotic defects of *Δcbf11* cells and its effect is not additive with ammonium supplementation. The *Δcbf11* mitotic defects can also be rescued, to varying degrees, by the introduction of a *tor1-D* temperature-sensitive allele, *tor1* deletion **(B)**, or deletion of *ssp2* **(C)**. Cells were grown to exponential phase in the indicated media and temperature, fixed, stained with DAPI and subjected to microscopy. Mean values ± SD from 3 independent experiments are shown. NA - not analysed. To determine statistical significance one-way Student’s t-test with Holm correction was applied.

Although known primarily as a Tor2/TORC1 inhibitor [59][60][61], rapamycin has a more general effect in *S. pombe*, inhibiting certain functions of Tor1/TORC2 as well [62][63]. Our next objective thus was to determine which branch of the TOR network was responsible for the rescue. To address this we first employed mutants of the non-essential *tor1* gene (TORC2). We found that the temperature-sensitive *tor1-D* allele [22] increased mitotic fidelity of the *Δcbf11* background when cells were grown at the semi-restrictive temperature of 34°C. Moreover, an even stronger rescue effect was achieved by introducing a complete deletion of *tor1* into *Δcbf11* cells (Fig. 7 B). It is important to note that the two TOR complexes are antagonistic and the inhibition or depletion of Tor1/TORC2 boosts the activity of Tor2/TORC1 [22][23].

Next, we focused on the role of the essential Tor2 kinase. First, we decreased Tor2/TORC1 kinase activity by combining the *Δcbf11* mutation with the temperature sensitive *tor2-S* allele [59]. We did not observe any significant change in the mitotic fidelity at semi-restrictive temperature compared to *Δcbf11* alone (Fig. S3). Then, to increase Tor2 activity we took advantage of the fact that the activity of Tor2/TORC1 is negatively regulated by the AMP-activated protein kinase (AMPK) complex [21]. Therefore, we deleted the *ssp2* gene, which encodes the AMPK catalytic subunit [64][65]. Indeed, the deletion of *ssp2* in *Δcbf11* cells led to a decrease in catastrophic mitotic events (Fig. 7 C). Collectively, these results strongly suggest that the nitrogen-dependent rescue of mitotic defects is mediated by Tor2/TORC1, and indicate a role for Tor2/TORC1 in ensuring successful progression through (closed) mitosis.

## DISCUSSION

The general availability and the particular quality of a nitrogen source have long been recognized as important factors regulating progression through the cell cycle. The absence of a nitrogen source (nitrogen starvation) or a shift to a less preferable nitrogen source (nitrogen stress) inhibit cell growth, and trigger quiescence or sexual differentiation, or accelerate the entry into mitosis, respectively [4][66]. However, we recently demonstrated that nitrogen also plays a role in mitotic fidelity in cells with perturbed lipid metabolism, with ammonium chloride being able to partially rescue catastrophic mitosis phenotypes [16][26]. Intriguingly, our current data revealed that this nitrogen-dependent rescue of mitosis is not accompanied by corrections of the aberrant FA and lipid composition, hinting at an indirect rescue mechanism. Indeed, we found that the rescue effect is of a more general nature, not limited to mutants in lipid metabolism. Moreover, we showed that nitrogen availability also affects the progression through mitosis, not just the G2/M transition (mitotic entry). While the mechanistic details remain to be elucidated, we demonstrated that the effect of nitrogen is mediated by the TOR regulatory network, namely the growth-promoting Tor2/TORC1 complex.

It was previously suggested that the mitotic defects of fission yeast cells with chemically or genetically perturbed lipid metabolism are caused by insufficient NE expansion due to a shortage of membrane precursors [6][17]. It is also possible that these mitotic defects could be related to altered mechanical properties of cell membranes that are important for spindle pole body integration and other crucial steps of the closed mitosis [53]. We have indeed found a number of aberrations in the composition of lipids in *Δcbf11* cells grown in the complex YES medium compared to WT. These aberrations included an overall lower content of FA, which are required for the production of new membranes. However, we did not observe any notable corrections in the mRNA levels of lipid metabolism genes (Fig. 3) or in composition of FA and lipids (Fig. 4, Fig. S1, Fig. S2) in *Δcbf11* cultures grown in an ammonium-supplemented YES medium, where mitotic defects are suppressed. Neither did we observe any significant improvement in NE expansion during mitosis in YES+ammonium (Fig. 2). These results are consistent with our recent findings that, in addition to any issues with the production of new membranes, mitotic fidelity in lipid metabolism mutants is affected by changes in centromeric chromatin structure and cohesin dynamics [18].

While we did not detect any nitrogen-dependent improvement in mitotic NE dynamics and FA and lipid composition composition, we found that ammonium supplementation clearly normalised the timing of mitotic progression, restoring the duration of individual mitotic phases close to their WT state (Fig. 5). Strikingly, we also found that the phenomenon of nitrogen-mediated rescue of mitotic fidelity is not limited to lipid metabolism-related problems, as mitotic defects in multiple unrelated ‘cut’ mutants could also be suppressed by ammonium supplementation (Fig. 6). Importantly, not all tested ‘cut’ mutants showed responsiveness to nitrogen availability, and among those which did, the magnitude of the rescue effect varied. Notably, the rescuable strains are typically mutants in genes involved in pre-anaphase processes, while all ammonium-insensitive ‘cut’ mutants are related to anaphase (securin/separase, APC/C) (Fig. 6). Thus, this specificity of the rescue effect may be dictated by the particular time during which the respective ‘cut’ genes perform their mitosis-related functions. Taken together, the ammonium-mediated rescue of mitotic defects is not limited to cells with perturbed lipid metabolism, and it seems to operate in cell cycle phase(s) prior to anaphase.

The TOR network is known to be one of the key cell-cycle regulators, promoting or inhibiting mitotic onset according to nutrient availability [19][67]. Interestingly, a previous report hinted that TOR may also have a role later in mitosis, as the viability of several temperature-sensitive separase and securin mutants (including those used in our study) was improved by treating cells with the TOR inhibitor rapamycin or by introducing the *tor2-S* mutation that impairs TORC1 kinase activity. However, the authors did not specifically test whether mitotic fidelity was also improved [22]. In our hands, the occurrence of mitotic defects in the *cut1-206* and *cut2-447* mutants was not suppressed by ammonium supplementation, a treatment we showed to have an impact similar to rapamycin treatment (Fig. 6). Neither did we observe any differences in mitotic fidelity between *Δcbf11* and *Δcbf11 tor2-S* cells (Fig. S3). It is therefore possible that any impact of TOR on the fidelity of anaphase events is regulated separately, by mechanism(s) different from those mediating the nitrogen-dependent rescue that we report here. Indeed, the authors themselves reported that the phenotype of the *tor2-S* mutant manifested clearly only in the peptone-containing YPD medium and not in the ammonium-containing EMM2, and they observed the *cut1*/*cut2* suppression under suboptimal nitrogen availability in YPD [22].

It was also reported that TOR is linked to growth phase-related changes in lipid metabolism. The switch between membrane phospholipid and storage triacylglycerol production is regulated by the lipin phosphatidic acid phosphatase [68]. In the budding yeast *Saccharomyces cerevisiae*, TOR controls lipin activity to ensure sufficient supply of phospholipids for new membrane production during active proliferation [69]. On the other hand, TOR/lipin-dependent overproduction of FA and endoplasmic reticulum membrane leads to mitotic defects and formation of micronuclei in mammals [70]. Nevertheless, we did not observe any major changes in phospholipid composition in *S. pombe* cells grown in ammonium supplemented YES medium compared to plain YES (Fig. S2).

We showed increased mitotic fidelity in *Δcbf11* cells when the stress-response branch of the TOR network was suppressed, either by ablation of Tor1/TORC2 or by boosting the activity of the pro-growth Tor2/TORC1 branch (Fig. 7). These data are in agreement with previous findings that Tor2/TORC1 inhibition mimics nitrogen starvation [71][72]. And vice versa, Tor2/TORC1 hyperactivation delays the response to nitrogen starvation [73], highlighting that the two branches of the TOR network are antagonistic and act in a negative feedback loop. Our results indicate that interventions providing cells with more, well-utilisable nitrogenous compounds (i.e. ammonium supplementation), or merely triggering internal signals mimicking the state of such nitrogen availability (i.e. boosting Tor2/TORC1 activity by ablating TORC2 or AMPK), make the mitotic defects in cells prone to catastrophic mitosis less severe. This could mean that the signalling of availability of a good nitrogen source is by itself more important for mitotic fidelity than the actual physical presence of the nutrients. However, the signalling of nutrient availability could in return affect the uptake and/or utilisation of nutrients by the cell. Additionally, it should be noted that ammonium-dependent improvements in mitotic dynamics manifested very early on during the mitosis of *Δcbf11* cells (Fig. 5). It is therefore also possible that the TOR network does not act on mitosis directly during the process of nuclear division, but rather earlier in the cell cycle helps establish conditions more favourable for smooth mitotic progression and increased fidelity further down the road.

In any case, the exact mechanism of the nitrogen-mediated rescue of mitotic fidelity remains to be characterised in detail, including the regulatory level(s) on which the observed effect is achieved. Since the TOR proteins are kinases, it is likely that the rescue is mediated, at least in part, at the post-translational level by phosphorylation of downstream effector proteins. These could be identified, for example, by screening of a knock-out library [74] combined with the *Δcbf11* mutation, as removal of these effectors should abolish the positive effects of nitrogen on mitotic fidelity and/or overall viability. Such studies could also explain the general character of the rescue, which occurs in a functionally diverse group of ‘cut’ mutants (Fig. 6).

TOR is linked to diabetes, cancer and other age-related diseases in humans [75]. Therefore, it would be interesting to see whether the novel mitotic role of TOR is conserved in metazoan cells, where both TORC1 and TORC2 complexes contain the same mTor kinase [76], and where open mitosis is employed. Currently, anti-cancer drugs targeting the human TOR are designed to inhibit the kinase activity. Unfortunately, such broad-impact TOR inhibition has so far shown limited applicability in cancer treatment (reviewed in [75][77]). Perhaps a more selective approach to targeting the TOR network may prove more suitable. In summary, we revealed a novel role of the TOR regulatory network as a factor influencing mitotic fidelity, thereby linking nutritional stimuli and metabolic state to the successful progression and completion of mitosis. We also revisited the previously identified importance of sufficient NE expansion for the fidelity of closed mitosis [6][17], and showed that it is not critical for the nitrogen-dependent mitotic rescue.

## DATA AVAILABILITY

RNA-seq data are available at ArrayExpress database (https://www.ebi.ac.uk/arrayexpress/) under the accession numbers: E-MTAB-13302 (WT, *Δcbf11*), E-MTAB-13303 (*Δmga2, Δmga2 Δcbf11*) and E-MTAB-13305 (WT, *cbf11DBM, Pcut6MUT,* and cerulenin treatment).

The scripts used for sequencing data processing and analyses are available at https://github.com/mprevorovsky/RNA-seq_ammonium (WT, *Δcbf11*), https://github.com/mprevorovsky/RNA-seq_CSL-DBM_cerulenin (WT, *cbf11DBM*, cerulenin treatment), and https://github.com/mprevorovsky/RNA-seq_mga2 (*Δmga2*, *Δmga2 Δcbf11*).

## ACKNOWLEDGEMENTS

We are very grateful to Phong Tran for his advice and material support for live-cell microscopy; Akshay Vishwanatha and Patrik Hohoš for the help with development of live-cell microscopy protocols; Adéla Kracíková and Kateřina Svobodová for excellent technical assistance.

The cut1-206, cut2-447, cut3-477, cut4-533, cut8-563, cut9-665, cut15-85, psm3-304, tor1-D and tor2-S strains were provided by NBRP, Japan. The smc6-x and nse3-R254E strains were provided by Jan Paleček.

Microscopy was performed in the Vinicna Microscopy Core Facility co-financed by the Czech-BioImaging large RI project LM2023050. Computational resources were supplied by the project “e-Infrastruktura CZ” (e-INFRA LM2018140) provided within the program Projects of Large Research, Development and Innovations Infrastructures.

The authors acknowledge Imaging Methods Core Facility at BIOCEV, institution supported by the MEYS CR (LM2023050 Czech-BioImaging) and for their support & assistance in this work.

The authors declare that they have no conflict of interest.

## SUPPLEMENTARY MATERIALS

**Figure S1.**
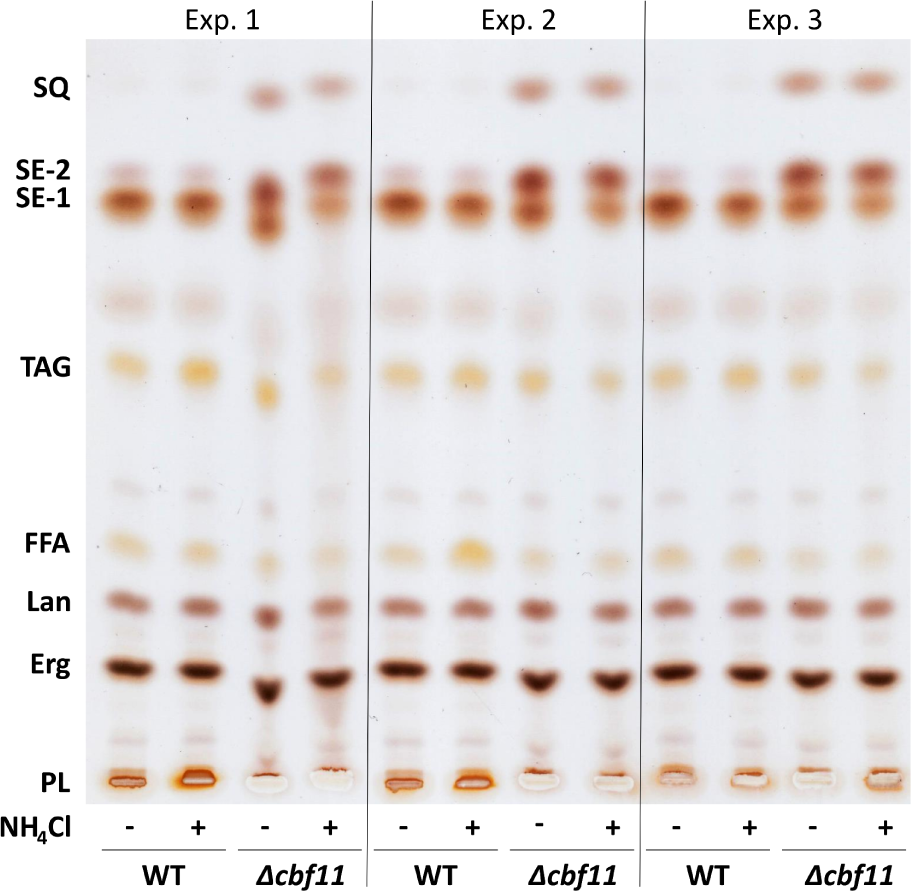
TLC analysis of neutral lipids. WT and *Δcbf11* cells were grown to exponential phase in the indicated media. Results from 3 independent experiments are shown. SQ - squalene; SE-1, SE-2 - sterylester species; TAG - triacylglycerol; FFA - free fatty acids; Lan - lanosterol; Erg - ergosterol; PL - phospholipids.

**Figure S2.**
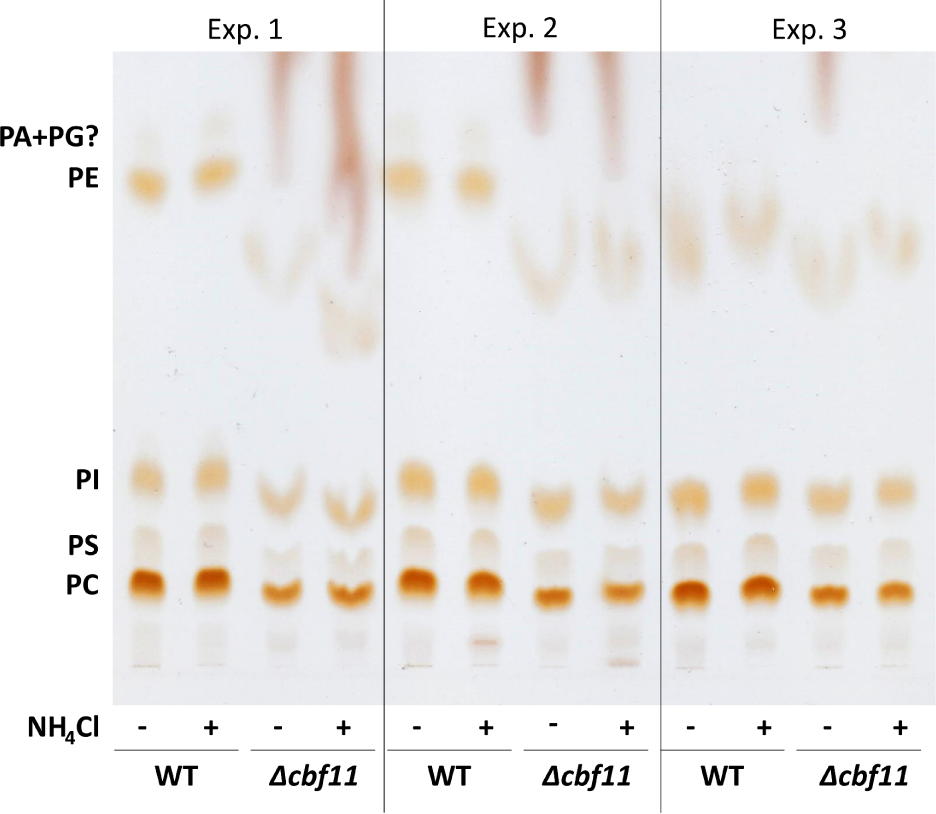
TLC analysis of phospholipids. WT and *Δcbf11* cells were grown to exponential phase in the indicated media. Results from 3 independent experiments are shown. PA+PG - phosphatidic acid + phosphatidylglycerol (presumed); PE - phosphatidylethanolamine; PI - phosphatidylinositol; PS - phosphatidylserine; PC - phosphatidylcholine.

**Figure S3.**
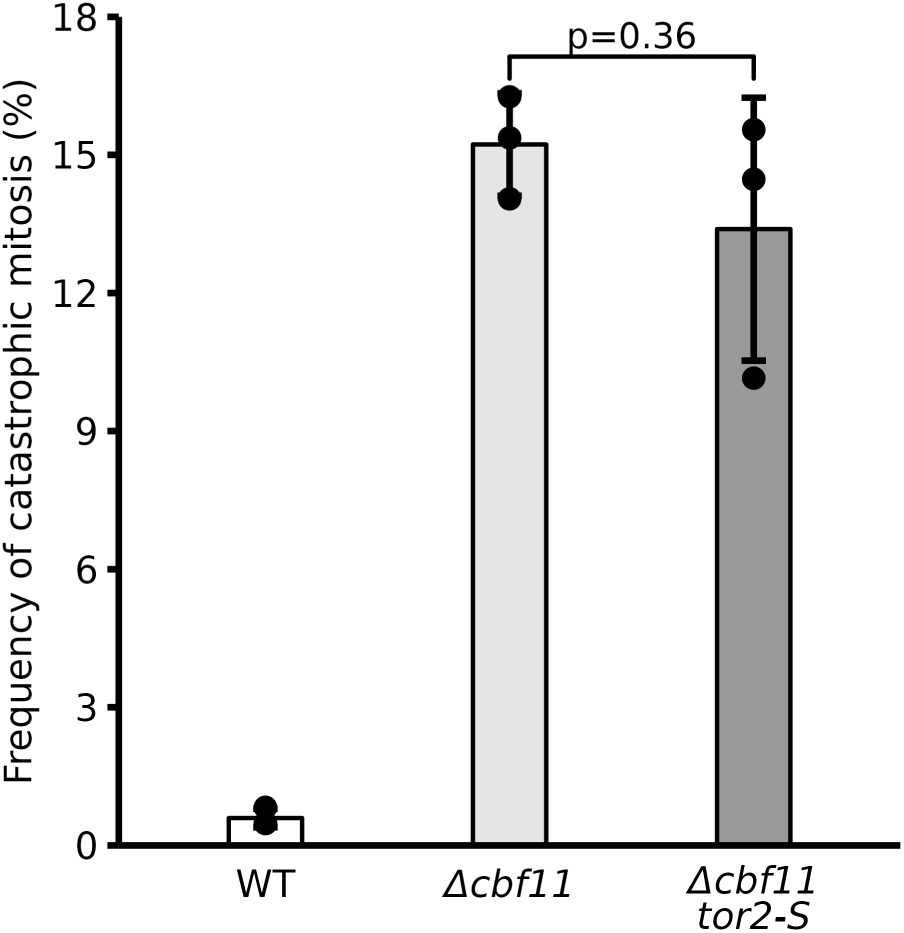
Suppression of Tor2/TORC1 does not affect mitotic fidelity. Mean values ± SD from 3 independent experiments are shown. To determine statistical significance two-way Student’s t-test was applied.

**Table S1.** Strains used in the study.

| ID | genotype | source |
| --- | --- | --- |
| JB22 | *h-* | lab stock |
| JB32 | *h+* | lab stock |
| MP44 | *h+ Δcbf11::kanR* | lab stock |
| Mcl316 | *h- cut11::GFP:ura4+ his3-D1 leu1-32 ura4-D18 ade6-M210* | Richard McIntosh |
| MP633 | *h+ ura4-D18* | lab stock |
| MP874 | *h+ cut11::GFP:ura4+ ura4-D18* | this study |
| M815 | *h+ Δmga2::natR* | lab stock |
| MP836 | *h+ Δmga2::natR Δcbf11::kanR* | lab stock |
| MP712 | *h+ cbf11DBM-ctap4* | lab stock |
| MP636 | *h- Pcut6MUT* | lab stock |
| FY11443 | *h- cut1-206* | NBRP |
| FY8129 | *h- leu1 cut2-447* | NBRP |
| MP794 | *h- cut2-447* | this study |
| FY7879 | *h+ cut3-477* | NBRP |
| FY9932 | *h- cut4-533* | NBRP |
| FY9497 | *h- cut8-563 leu1* | NBRP |
| MP798 | *h- cut8-563* | this study |
| FY9685 | *h+ cut9-665* | NBRP |
| FY17627 | *h- cut15-85 leu1* | NBRP |
| MP800 | *h+ cut15-85* | this study |
| FY20148 | *h+ psm3-304* | NBRP |
| 5C7 | *h- smc6-x leu1-32 ade6-704 ura4-d18* | Jan Paleček |
| MP802 | *h+ smc6-x* | this study |
| 3H5 | *h+ nse3-R254E leu1-32 ade6-704 ura4-d18* | Jan Paleček |
| MP806 | *h- nse3-R254E* | this study |
| TP 3640 | *h+ mCherry-atb2-HygR hht2-GFP:Ura4+ ade6-21? leu1-32 ura4-D18* | Phong Tran |
| MP807 | *h+ mCherry-atb2-HygR hht2-GFP:Ura4+ ura4-D18* | lab stock |
| MP416 | *h+ Δtor1:kanR* | Charalampos Rallis |
| MP973 | *Δtor1::ura4 Δcbf11::natR* | this study |
| FY25826 | *tor1-L2045D:hygR* (“*tor1-D*”) | NBRP |
| MP976 | *h- tor1-L2045D:hygR Δcbf11::natR* | this study |
| FY21141 | *tor2-L2048S* (“*tor2-S*”) | NBRP |
| MP859 | *h- tor2-L2048S Δcbf11::natR* | this study |
| MP635 | *h+ Δssp2::natR* | lab stock |
| MP855 | *h+ Δssp2::natR Δcbf11::hygR* | this study |

**Table S2.**
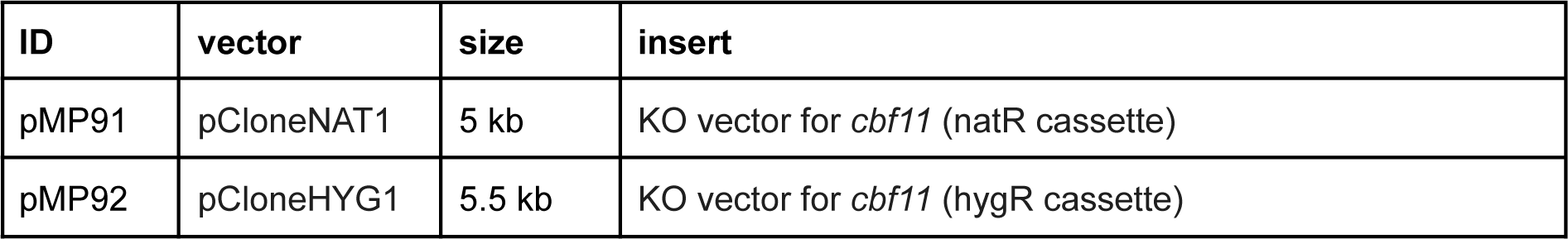
Plasmids used in the study.

**Table S3.**
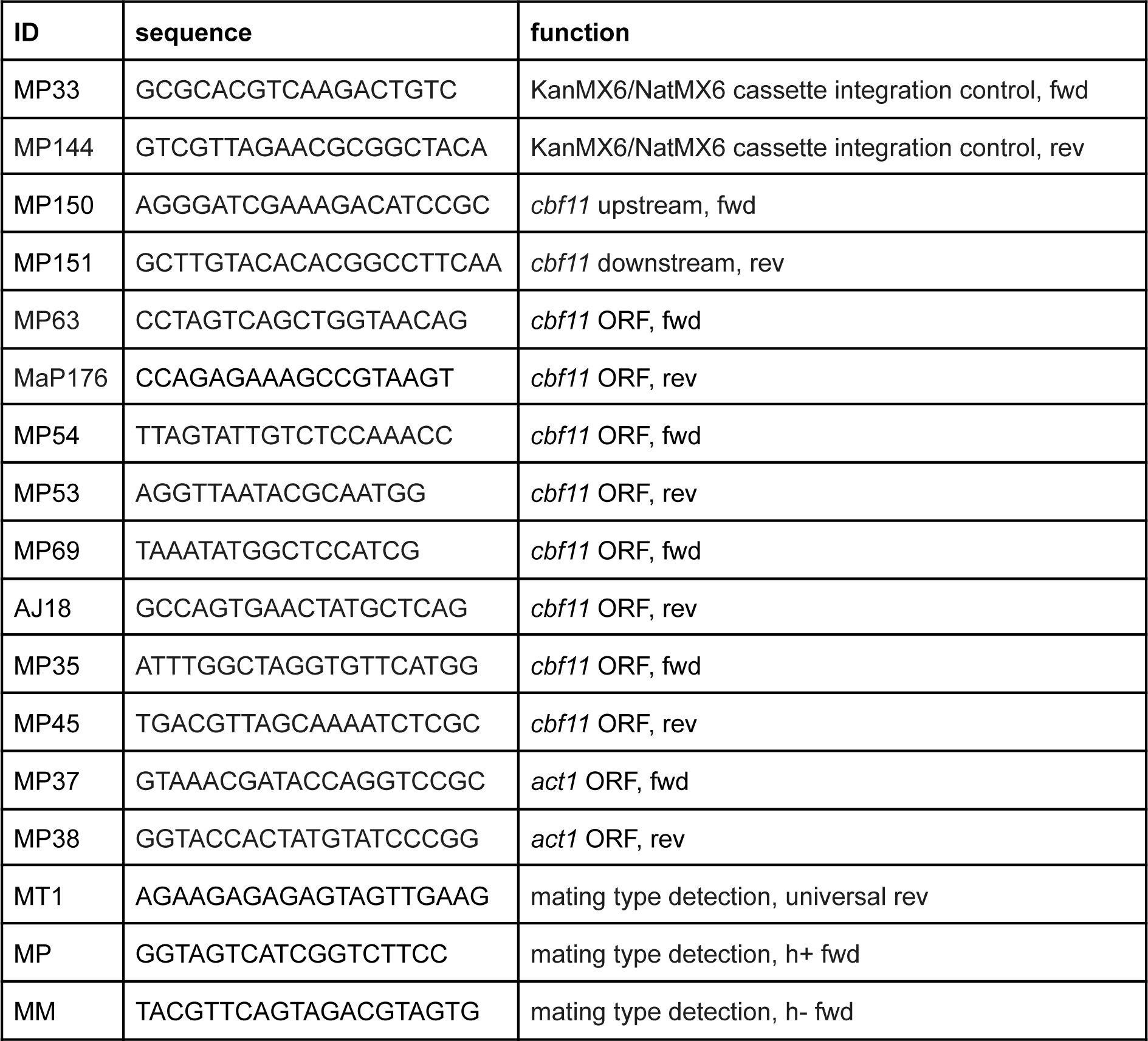
Oligonucleotides used in the study.

## REFERENCES

[1] Saeki, T., Ouchi, M., & Ouchi, T. (2009). Physiological and Oncogenic Aurora-A Pathway. International Journal of Biological Sciences, 5(7), 758–762. 10.7150/IJBS.5.758

[2] Hayles, J., Wood, V., Jeffery, L., Hoe, K. L., Kim, D. U., Park, H. O., Salas-Pino, S., Heichinger, C., & Nurse, P. (2013). A genome-wide resource of cell cycle and cell shape genes of fission yeast. Open Biology, 3(5), 130053. 10.1098/RSOB.130053

[3] Bähler, J. (2005). Cell-cycle control of gene expression in budding and fission yeast. In Annual Review of Genetics (Vol. 39, pp. 69–94). Annual Reviews. 10.1146/annurev.genet.39.110304.095808

[4] Fantes, P., & Nurse, P. (1977). Control of cell size at division in fission yeast by a growth-modulated size control over nuclear division. Experimental Cell Research, 107(2), 377–386. 10.1016/0014-4827(77)90359-7

[5] Boettcher, B., & Barral, Y. (2013). The cell biology of open and closed mitosis. Nucleus (United States), 4(3), 160–165. 10.4161/nucl.24676

[6] Yam, C., He, Y., Zhang, D., Chiam, K. H., & Oliferenko, S. (2011). Divergent strategies for controlling the nuclear membrane satisfy geometric constraints during nuclear division. Current Biology, 21(15), 1314–1319. 10.1016/j.cub.2011.06.052

[7] Uemura, T., & Yanagida, M. (1984). Isolation of type I and II DNA topoisomerase mutants from fission yeast: single and double mutants show different phenotypes in cell growth and chromatin organization. The EMBO Journal, 3(8), 1737–1744. 10.1002/j.1460-2075.1984.tb02040.x

[8] Hirano, T., Funahashi, S., Uemura, T., & Yanagida, M. (1986). Isolation and characterization of Schizosaccharomyces pombe cut mutants that block nuclear division but not cytokinesis. The EMBO Journal, 5(11), 2973–2979. 10.1002/j.1460-2075.1986.tb04594.x

[9] Saitoh, S., Takahashi, K., Nabeshima, K., Yamashita, Y., Nakaseko, Y., Hirata, A., & Yanagida, M. (1996). Aberrant mitosis in fission yeast mutants defective in fatty acid synthetase and acetyl CoA carboxylase. The Journal of Cell Biology, 134(4), 949. 10.1083/JCB.134.4.949

[10] Yukawa, M., Teratani, Y., & Toda, T. (2021). Escape from mitotic catastrophe by actin-dependent nuclear displacement in fission yeast. IScience, 24(1), 102031. 10.1016/J.ISCI.2020.102031

[11] Uzawa, S., Samejima, I., Hirano, T., Tanaka, K., & Yanagida, M. (1990). The fission yeast cut1+ gene regulates spindle pole body duplication and has homology to the budding yeast ESP1 gene. Cell, 62(5), 913–925. 10.1016/0092-8674(90)90266-H

[12] Saka, Y., Sutani, T., Yamashita, Y., Saitoh, S., Takeuchi, M., Nakaseko, Y., & Yanagida, M. (1994). Fission yeast cut3 and cut14, members of a ubiquitous protein family, are required for chromosome condensation and segregation in mitosis. EMBO Journal, 13(20), 4938–4952. 10.1002/j.1460-2075.1994.tb06821.x

[13] Yanagida, M., Yamashita, Y. M., Tatebe, H., Ishii, K., Kumada, K., & Nakaseko, Y. (1999). Control of metaphase-anaphase progression by proteolysis: cyclosome function regulated by the protein kinase A pathway, ubiquitination and localization. Philosophical Transactions of the Royal Society of London. Series B: Biological Sciences, 354(1389), 1559–1570. 10.1098/RSTB.1999.0499

[14] Yanagida, M. (1998). Fission yeast cut mutations revisited: Control of anaphase. Trends in Cell Biology, 8(4), 144–149. 10.1016/S0962-8924(98)01236-7

[15] Zach, R., & Převorovský, M. (2018). The phenomenon of lipid metabolism “cut” mutants. Yeast, 35(12), 631–637. 10.1002/yea.3358

[16] Převorovský, M., Oravcová, M., Tvarůžková, J., Zach, R., Folk, P., Půta, F., & Bähler, J. (2015). Fission yeast CSL transcription factors: Mapping their target genes and biological roles. PLoS ONE, 10(9), 1–23. 10.1371/journal.pone.0137820

[17] Takemoto, A., Kawashima, S. A., Li, J. J., Jeffery, L., Yamatsugu, K., Elemento, O., & Nurse, P. (2016). Nuclear envelope expansion is crucial for proper chromosomal segregation during a closed mitosis. Journal of Cell Science, 129(6), 1250–1259. 10.1242/jcs.181560

[18] Vishwanatha, A., Princová, J., Hohoš, P., Zach, R., & Převorovský, M. (2023). Altered cohesin dynamics and H3K9 modifications contribute to mitotic defects in the cbf11Δ lipid metabolism mutant. Journal of Cell Science, 136(11). 10.1242/jcs.261265

[19] Petersen, J., & Nurse, P. (2007). TOR signalling regulates mitotic commitment through the stress MAP kinase pathway and the Polo and Cdc2 kinases. Nature Cell Biology, 9(11), 1263–1272. 10.1038/ncb1646

[20] Yanagida, M., Ikai, N., Shimanuki, M., & Sajiki, K. (2011). Nutrient limitations alter cell division control and chromosome segregation through growth-related kinases and phosphatases. In Philosophical Transactions of the Royal Society B: Biological Sciences (Vol. 366, Issue 1584, pp. 3508–3520). The Royal Society. 10.1098/rstb.2011.0124

[21] Davie, E., Forte, G. M. A., & Petersen, J. (2015). Nitrogen regulates AMPK to control TORC1 signaling. Current Biology, 25(4), 445–454. 10.1016/j.cub.2014.12.034

[22] Ikai, N., Nakazawa, N., Hayashi, T., & Yanagida, M. (2011). The reverse, but coordinated, roles of Tor2 (TORC1) and Tor1 (TORC2) kinases for growth, cell cycle and separase-mediated mitosis in Schizosaccharomyces pombe. Open Biology, 1(NOVEMBER). 10.1098/rsob.110007

[23] Schonbrun, M., Laor, D., López-Maury, L., Bähler, J., Kupiec, M., & Weisman, R. (2009). TOR Complex 2 Controls Gene Silencing, Telomere Length Maintenance, and Survival under DNA-Damaging Conditions. Molecular and Cellular Biology, 29(16), 4584–4594. 10.1128/mcb.01879-08

[24] Petersen, J., & Russell, P. (2016). Growth and the environment of Schizosaccharomyces pombe. Cold Spring Harbor Protocols, 2016(3), 210–226. 10.1101/pdb.top079764

[25] Carlson, C. R., Grallert, B., Stokke, T., & Boye, E. (1999). Regulation of the start of DNA replication in Schizosaccharomyces pombe. Journal of Cell Science, 112(6), 939–946. 10.1242/JCS.112.6.939

[26] Zach, R., Tvarůžková, J., Schätz, M., Tupa, O., Grallert, B., & Převorovský, M. (2018). Mitotic defects in fission yeast lipid metabolism ‘cut’ mutants are suppressed by ammonium chloride. FEMS Yeast Research, 18(6), 1–7. 10.1093/femsyr/foy064

[27] Sabatinos, S. A., & Forsburg, S. L. (2010). Molecular genetics of Schizosaccharomyces pombe. Methods in Enzymology, 470(C), 759–795. 10.1016/S0076-6879(10)70032-X

[28] Gregan, J., Rabitsch, P. K., Rumpf, C., Novatchkova, M., Schleiffer, A., & Nasmyth, K. (2006). High-throughput knockout screen in fission yeast. Nature Protocols, 1(5), 2457. 10.1038/NPROT.2006.385

[29] Costa, J., Fu, C., Syrovatkina, V., & Tran, P. T. (2013). Imaging individual spindle microtubule dynamics in fission yeast. In Methods in Cell Biology (Vol. 115, pp. 385–394). Academic Press. 10.1016/B978-0-12-407757-7.00024-4

[30] Schindelin, J., Arganda-Carreras, I., Frise, E., Kaynig, V., Longair, M., Pietzsch, T., Preibisch, S., Rueden, C., Saalfeld, S., Schmid, B., Tinevez, J. Y., White, D. J., Hartenstein, V., Eliceiri, K., Tomancak, P., & Cardona, A. (2012). Fiji: An open-source platform for biological-image analysis. Nature Methods, 9(7), 676–682. 10.1038/nmeth.2019

[31] Miura, K., Cordelières, F. P., & Klemm, A. H. (2020). Bleach correction ImageJ plugin for compensating the photobleaching of time-lapse sequences. F1000Research 2020 9:1494, 9, 1494. 10.12688/f1000research.27171.1

[32] Lowe, D. G. (2004). Distinctive image features from scale-invariant keypoints. International Journal of Computer Vision, 60(2), 91–110. 10.1023/B:VISI.0000029664.99615.94

[33] Lyne, R., Burns, G., Mata, J., Penkett, C. J., Rustici, G., Chen, D., Langford, C., Vetrie, D., & Bähler, J. (2003). Whole-genome microarrays of fission yeast: Characteristics, accuracy, reproducibility, and processing of array data. BMC Genomics, 4(1), 1–15. 10.1186/1471-2164-4-27/FIGURES/7

[34] Wood, V., Gwilliam, R., Rajandream, M. A., Lyne, M., Lyne, R., Stewart, A., Sgouros, J., Peat, N., Hayles, J., Baker, S., Basham, D., Bowman, S., Brooks, K., Brown, D., Brown, S., Chillingworth, T., Churcher, C., Collins, M., Connor, R., … Nurse, P. (2002). The genome sequence of Schizosaccharomyces pombe. Nature 2002 415:6874, 415(6874), 871–880. 10.1038/nature724

[35] Lock, A., Rutherford, K., Harris, M. A., Hayles, J., Oliver, S. G., Bähler, J., & Wood, V. (2019). PomBase 2018: user-driven reimplementation of the fission yeast database provides rapid and intuitive access to diverse, interconnected information. Nucleic Acids Research, 47(D1), D821–D827. 10.1093/NAR/GKY961

[36] Andrews S. (2010). FastQC: a quality control tool for high throughput sequence data. Available online at: http://www.bioinformatics.babraham.ac.uk/projects/fastqc

[37] Bolger, A. M., Lohse, M., & Usadel, B. (2014). Trimmomatic: a flexible trimmer for Illumina sequence data. Bioinformatics, 30(15), 2114–2120. 10.1093/BIOINFORMATICS/BTU170

[38] Kim, D., Langmead, B., & Salzberg, S. L. (2015). HISAT: a fast spliced aligner with low memory requirements. Nature Methods 2015 12:4, 12(4), 357–360. 10.1038/nmeth.3317

[39] Li, H., Handsaker, B., Wysoker, A., Fennell, T., Ruan, J., Homer, N., Marth, G., Abecasis, G., & Durbin, R. (2009). The Sequence Alignment/Map format and SAMtools. Bioinformatics, 25(16), 2078–2079. 10.1093/BIOINFORMATICS/BTP352

[40] Ramírez, F., Ryan, D. P., Grüning, B., Bhardwaj, V., Kilpert, F., Richter, A. S., Heyne, S., Dündar, F., & Manke, T. (2016). deepTools2: a next generation web server for deep-sequencing data analysis. Nucleic Acids Research, 44(W1), W160–W165. 10.1093/NAR/GKW257

[41] Robinson, J. T., Thorvaldsdóttir, H., Winckler, W., Guttman, M., Lander, E. S., Getz, G., & Mesirov, J. P. (2011). Integrative genomics viewer. Nature Biotechnology 2011 29:1, 29(1), 24–26. 10.1038/nbt.1754

[42] Lawrence, M., Huber, W., Pagès, H., Aboyoun, P., Carlson, M., Gentleman, R., Morgan, M. T., & Carey, V. J. (2013). Software for Computing and Annotating Genomic Ranges. PLOS Computational Biology, 9(8), e1003118. 10.1371/JOURNAL.PCBI.1003118

[43] R Core Team (2023). R: A Language and Environment for Statistical Computing. R Foundation for Statistical Computing, Vienna, Austria. Available online at: https://www.R-project.org

[44] Gentleman, R. C., Carey, V. J., Bates, D. M., Bolstad, B., Dettling, M., Dudoit, S., Ellis, B., Gautier, L., Ge, Y., Gentry, J., Hornik, K., Hothorn, T., Huber, W., Iacus, S., Irizarry, R., Leisch, F., Li, C., Maechler, M., Rossini, A. J., … Zhang, J. (2004). Bioconductor: open software development for computational biology and bioinformatics. Genome Biology 2004 5:10, 5(10), 1–16. 10.1186/GB-2004-5-10-R80

[45] Risso, D., Ngai, J., Speed, T. P., & Dudoit, S. (2014). Normalization of RNA-seq data using factor analysis of control genes or samples. Nature Biotechnology, 32(9), 896–902. 10.1038/nbt.2931

[46] Love, M. I., Huber, W., & Anders, S. (2014). Moderated estimation of fold change and dispersion for RNA-seq data with DESeq2. Genome Biology, 15(12), 1–21. 10.1186/S13059-014-0550-8/FIGURES/9

[47] Garaiova, M.; Mietkiewska, E.; Weselake, R. J.; Holic, R. (2017) Metabolic Engineering of Schizosaccharomyces Pombe to Produce Punicic Acid, a Conjugated Fatty Acid with Nutraceutic Properties. Appl Microbiol Biotechnol 101, 7913–7922. 10.1007/s00253-017-8498-8

[48] Iwabuchi, M.; Kohno-Murase, J.; Imamura, J. (2003) Delta 12-Oleate Desaturase-Related Enzymes Associated with Formation of Conjugated Trans-Delta 11, Cis-Delta 13 Double Bonds. J Biol Chem, 278 (7), 4603–4610. 10.1074/jbc.M210748200

[49] Mietkiewska, E.; Siloto, R. M.; Dewald, J.; Shah, S.; Brindley, D. N.; Weselake, R. J. (2011) Lipins from Plants Are Phosphatidate Phosphatases That Restore Lipid Synthesis in a Pah1Delta Mutant Strain of Saccharomyces Cerevisiae. FEBS J, 278 (5), 764–775. 10.1111/j.1742-4658.2010.07995.x.

[50] Spanova M, Czabany T, Zellnig G, Leitner E, Hapala I, Daum G (2010) Effect of lipid particle biogenesis on the subcellular distribution of squalene in the yeast Saccharomyces cerevisiae J Biol Chem 285:6127–6133 10.1074/jbc.M109.074229

[51] Garaiová, M.; Zambojová, V.; Šimová, Z.; Griač, P.; Hapala, I. (2014) Squalene Epoxidase as a Target for Manipulation of Squalene Levels in the Yeast Saccharomyces Cerevisiae. FEMS Yeast Res., 14, 310–323, 10.1111/1567-1364.12107

[52] Garner, K.; Hunt, A.N.; Koster, G.; Somerharju, P.; Groves, E.; Li, M.; Raghu, P.; Holic, R.; Cockcroft, S. (2012) Phosphatidylinositol Transfer Protein, Cytoplasmic 1 (PITPNC1) Binds and Transfers Phosphatidic Acid. J. Biol. Chem., 287, 32263–32276, 10.1074/jbc.M112.375840.

[53] West, R. R., Vaisberg, E. V., Ding, R., Nurse, P., & Richard McIntosh, J. (1998). cut11+: A gene required for cell cycle-dependent spindle pole body anchoring in the nuclear envelope and bipolar spindle formation in Schizosaccharomyces pombe. Molecular Biology of the Cell, 9(10), 2839–2855. 10.1091/mbc.9.10.2839

[54] Princová, J., Salat-Canela, C., Daněk, P., Marešová, A., de Cubas, L., Bähler, J., Ayté, J., Hidalgo, E., & Převorovský, M. (2023). Perturbed fatty-acid metabolism is linked to localized chromatin hyperacetylation, increased stress-response gene expression and resistance to oxidative stress. PLoS Genetics, 19(1). 10.1371/JOURNAL.PGEN.1010582

[55] Burr, R., Stewart, E. V., Shao, W., Zhao, S., Hannibal-Bach, H. K., Ejsing, C. S., & Espenshade, P. J. (2016). Mga2 Transcription Factor Regulates an Oxygen-responsive Lipid Homeostasis Pathway in Fission Yeast. Journal of Biological Chemistry, 291(23), 12171–12183. 10.1074/JBC.M116.723650

[56] Převorovský, M., Oravcová, M., Zach, R., Jordáková, A., Bähler, J., Půta, F., & Folk, P. (2016). CSL protein regulates transcription of genes required to prevent catastrophic mitosis in fission yeast. Cell Cycle, 15(22), 3082–3093. 10.1080/15384101.2016.1235100

[57] Burr, R., Stewart, E. V., & Espenshade, P. J. (2017). Coordinate regulation of yeast sterol regulatory elementbinding protein (SREBP) and Mga2 transcription factors. Journal of Biological Chemistry, 292(13), 5311–5324. 10.1074/jbc.M117.778209

[58] Heitman, J., Movva, N. R., & Hall, M. N. (1991). Targets for cell cycle arrest by the immunosuppressant rapamycin in yeast. Science, 253(5022), 905–909. 10.1126/science.1715094

[59] Hayashi, T., Hatanaka, M., Nagao, K., Nakaseko, Y., Kanoh, J., Kokubu, A., Ebe, M., & Yanagida, M. (2007). Rapamycin sensitivity of the Schizosaccharomyces pombe tor2 mutant and organization of two highly phosphorylated TOR complexes by specific and common subunits. Genes to Cells, 12(12), 1357–1370. 10.1111/j.1365-2443.2007.01141.x

[60] Nakashima, A., Sato, T., & Tamanoi, F. (2010). Fission yeast TORC1 regulates phosphorylation of ribosomal S6 proteins in response to nutrients and its activity is inhibited by rapamycin. Journal of Cell Science, 123(5), 777–786. 10.1242/jcs.060319

[61] Takahara, T., & Maeda, T. (2012). TORC1 of fission yeast is rapamycin-sensitive. Genes to Cells, 17(8), 698–708. 10.1111/j.1365-2443.2012.01618.x

[62] Weisman, R., Finkelstein, S., & Choder, M. (2001). Rapamycin Blocks Sexual Development in Fission Yeast through Inhibition of the Cellular Function of an FKBP12 Homolog. Journal of Biological Chemistry, 276(27), 24736–24742. 10.1074/jbc.M102090200

[63] Weisman, R., Roitburg, I., Nahari, T., & Kupiec, M. (2005). Regulation of leucine uptake by tor1+ in Schizosaccharomyces pombe is sensitive to rapamycin. Genetics, 169(2), 539–550. 10.1534/genetics.104.034983

[64] Matsuzawa, T., Fujita, Y., Tohda, H., & Takegawa, K. (2012). Snf1-like protein kinase ssp2 regulates glucose derepression in schizosaccharomyces pombe. Eukaryotic Cell, 11(2), 159–167. 10.1128/EC.05268-11

[65] Valbuena, N., & Moreno, S. (2012). AMPK phosphorylation by Ssp1 is required for proper sexual differentiation in fission yeast. Journal of Cell Science, 125(11), 2655–2664. 10.1242/jcs.098533

[66] Egel, R., & Egel-Mitani, M. (1974). Premeiotic DNA synthesis in fission yeast. Experimental Cell Research, 88(1), 127–134. 10.1016/0014-4827(74)90626-0

[67] Uritani, M., Hidaka, H., Hotta, Y., Ueno, M., Ushimaru, T., & Toda, T. (2006). Fission yeast Tor2 links nitrogen signals to cell proliferation and acts downstream of the Rheb GTPase. Genes to Cells, 11(12), 1367–1379. 10.1111/J.1365-2443.2006.01025.X

[68] Santos-Rosa, H., Leung, J., Grimsey, N., Peak-Chew, S., & Siniossoglou, S. (2005). The yeast lipin Smp2 couples phospholipid biosynthesis to nuclear membrane growth. EMBO Journal, 24(11), 1931–1941. 10.1038/sj.emboj.7600672

[69] Dubots, E., Cottier, S., Péli-Gulli, M. P., Jaquenoud, M., Bontron, S., Schneiter, R., & De Virgilio, C. (2014). TORC1 regulates Pah1 phosphatidate phosphatase activity via the Nem1/Spo7 protein phosphatase complex. PLoS ONE, 9(8), e104194. 10.1371/journal.pone.0104194

[70] Merta, H., Carrasquillo Rodríguez, J. W., Anjur-Dietrich, M. I., Vitale, T., Granade, M. E., Harris, T. E., Needleman, D. J., & Bahmanyar, S. (2021). Cell cycle regulation of ER membrane biogenesis protects against chromosome missegregation. Developmental Cell, 56(24), 3364–3379.e10. 10.1016/j.devcel.2021.11.009

[71] Matsuo, T., Otsubo, Y., Urano, J., Tamanoi, F., & Yamamoto, M. (2007). Loss of the TOR Kinase Tor2 Mimics Nitrogen Starvation and Activates the Sexual Development Pathway in Fission Yeast. Molecular and Cellular Biology, 27(8), 3154–3164. 10.1128/mcb.01039-06

[72] Weisman, R., Roitburg, I., Schonbrun, M., Harari, R., & Kupiec, M. (2007). Opposite effects of Tor1 and Tor2 on nitrogen starvation responses in fission yeast. Genetics, 175(3), 1153–1162. 10.1534/genetics.106.064170

[73] Urano, J., Sato, T., Matsuo, T., Otsubo, Y., Yamamoto, M., & Tamanoi, F. (2007). Point mutations in TOR confer Rheb-independent growth in fission yeast and nutrient-independent mammalian TOR signaling in mammalian cells. Proceedings of the National Academy of Sciences of the United States of America, 104(9), 3514. 10.1073/PNAS.0608510104

[74] Kim, D. U., Hayles, J., Kim, D., Wood, V., Park, H. O., Won, M., Yoo, H. S., Duhig, T., Nam, M., Palmer, G., Han, S., Jeffery, L., Baek, S. T., Lee, H., Shim, Y. S., Lee, M., Kim, L., Heo, K. S., Noh, E. J., … Hoe, K. L. (2010). Analysis of a genome-wide set of gene deletions in the fission yeast Schizosaccharomyces pombe. Nature Biotechnology, 28(6), 617–623. 10.1038/nbt.1628

[75] Boutouja, F., Stiehm, C. M., & Platta, H. W. (2019). mTOR: A cellular regulator interface in health and disease. Cells (Vol. 8, Issue 1). 10.3390/cells8010018

[76] Aspuria, P. J., Sato, T., & Tamanoi, F. (2007). The TSC/Rheb/TOR signaling pathway in fission yeast and mammalian cells: Temperature sensitive and constitutive active mutants of TOR. In Cell Cycle (Vol. 6, Issue 14, pp. 1692–1695). Taylor & Francis. 10.4161/cc.6.14.4478

[77] Zheng, Y., & Jiang, Y. (2015). mTOR inhibitors at a glance. Molecular and Cellular Pharmacology, 7(2), 15–20. 10.4255/mcpharmacol.15.02

